# ^31^P Transversal Relaxation Times and Metabolite Concentrations in the Human Brain at 9.4T

**DOI:** 10.1101/2021.12.28.474330

**Authors:** Johanna Dorst, Tamas Borbath, Loreen Ruhm, Anke Henning

**Affiliations:** High-Field MR Center, Max Planck Institute for Biological Cybernetics, Tübingen, Germany; IMPRS for Cognitive and Systems Neuroscience, University of Tübingen, Tübingen, Germany; Faculty of Science, University of Tübingen, Tübingen, Germany; Advanced Imaging Research Center, UT Southwestern Medical Center, Dallas, TX, United States

**Keywords:** phosphorus, ultrahigh field, 9.4T, STEAM, T2, healthy human brain, J-evolution

## Abstract

A method to estimate phosphorus (^31^P) transversal relaxation times (T_2_) of coupled spin systems is demonstrated. Additionally, intracellular and extracellular pH (pH_ext_, pH_int_) and relaxation corrected metabolite concentrations are reported. Echo time (TE) series of ^31^P metabolite spectra were acquired using STEAM localization. Spectra were fitted using LCModel with accurately modeled Vespa basis sets accounting for J-evolution of the coupled spin systems. T_2_s were estimated by fitting a single exponential two-parameter model across the TE series. Fitted inorganic phosphate frequencies were used to calculate pH, and relaxation times were used to determine the brain metabolite concentrations. The method was demonstrated in the healthy human brain at a field strength of 9.4T. T_2_ relaxation times of ATP and NAD are the shortest between 8 ms and 20 ms, followed by T_2_s of inorganic phosphate between 25 ms and 50 ms, and PCr with a T_2_ of 100 ms. Phosphomonoesters and –diesters have the longest T_2_s of about 130 ms. Measured T_2_s are comparable to literature values and fit in a decreasing trend with increasing field strengths. Calculated pHs and metabolite concentrations are also comparable to literature values.

## Introduction

In vivo phosphorus MR Spectroscopy (^31^P-MRS) provides insight into energy and phospholipid membrane metabolism noninvasively^1^. It also offers the possibility for intracellular and extracellular pH measurements, or the determination of intracellular Mg^2+^ concentration^1,2^. Thereby, this technique is used for different applications, such as pH mapping in cancer^3,4^, measurement of altered phospholipid and energy-related metabolites in schizophrenia^5^, or measurement of free cytosolic magnesium in neurodegenerative disorders or migraine^6,7^.

So far, ^31^P MRS is rarely used for clinical applications due to the low intrinsic sensitivity of ^31^P, and therefore its low spatial and temporal resolution. Since the signal to noise ratio (SNR) is dependent on the static magnetic field B_0_, a sensitivity gain and spectral improvement can be obtained at higher field strengths allowing shorter acquisition times and higher spatial resolution^8^. In addition to clinically acceptable measurement times, the comparison of MRS results between patients, different scanners or acquisition methods is mandatory for the diagnosis of diseases. Such a comparison requires reliable quantitative evaluation of metabolite concentrations and accurate knowledge about relaxation times is a prerequisite to achieving this. Relaxation times may also be of interest for characterizing molecular dynamics^9^. For setting up measurement protocols and for optimizing respective pulse sequences concerning repetition time, echo time, and free precession times, knowledge on relaxation times is essential as well. Since relaxation times are field strength dependent, they need to be measured at every field strength^10,11^.

Several studies show an increase in T_1_ and decrease in T_2_ relaxation times in the human brain for protons with increasing field strengths^1,10,12^. In contrast, a decrease in T_1_ with increasing field strength was observed for ^31^P metabolites in the human brain^8,9^. For human in vivo T_2_ relaxation times, no coherent trend with magnetic field strength can be extracted from literature^11,13–16^. Especially reports on transversal relaxation times of ATP largely vary^11,13–15^. These discrepancies originate from different acquisition and processing techniques. In some of the studies, the homonuclear scalar coupling of the ATP molecule, which leads to TE-dependent phase and amplitude modulations of the signals, was not considered^13,17,18^. Therefore, these studies report underestimated T_2_ relaxation times^14,15^. Adjusted acquisition methods using selective refocusing pulses or homonuclear decoupling allow the accurate determination of T_2_ relaxation times^11,13,19^.

None of the studies measuring ^31^P T_2_ relaxation times in the human brain used the approach to consider scalar coupling in the fit procedure. In a frequency domain fitting approach like LCModel, spectra are approximated as a linear combination of model spectra^20^. Spectral simulation of the model spectra, for instance, performed with Vespa or MARSS, are specific for each sequence and timing and fully consider J-evolution^21,22^. Using this approach, no selective refocusing pulses or homonuclear decoupling is needed in the acquisition sequence. Since ^31^P T_2_ relaxation times are expected to be very short at an ultrahigh field strength (UHF) and are typically estimated by observing the exponential signal decay in echo time (TE) series spectra, a localization sequence that allows short TEs should be used. Frequency-selective refocusing or homonuclear decoupling used in previous studies lead to SAR-related concerns at UHF11,13,16,18,19,23.

The primary goal of this study was to demonstrate a method to measure ^31^P T_2_ relaxation times of J-coupled metabolites without frequency-selective refocusing or homonuclear decoupling, as was done in previous studies^24,25^. Therefore, TE series spectra over a broad frequency range were acquired with a stimulated echo acquisition mode (STEAM) localization sequence optimized in a previous study^24^ entailing a high SNR per unit of time, which is a critical factor for T_2_ measurements. J-modulation of the ^31^P metabolites was then considered in the fitting routine using Vespa in combination with LCModel^20,21^. This approach to estimate ^31^P T_2_ relaxation times was demonstrated in the human brain at a field strength of 9.4T. Additionally, pH values as well as estimated tissue concentrations of ^31^P metabolites after applying relaxation corrections are reported. T_2_ values and metabolite concentrations are compared to literature data, and factors affecting their accuracy and comparability are discussed.

## Experimental

### Study design

All data were acquired at a 9.4 T whole-body MRI scanner (Siemens, Erlangen, Germany) using a home-built double-tuned 20-loop ^31^P/^1^H head array^26^. To increase B_1_ ^+^ for ^31^P single-voxel spectroscopy in the occipital lobe, the entire RF power was applied only to the three bottom surface coil elements using an unbalanced three-way Wilkinson power splitter, as described recently^24^.

All measurements with volunteers were in accordance with the local research ethics guidelines, and written informed consent was obtained from all volunteers before the examination. Twelve healthy volunteers participated in the study (7 females, 5 males, age 27 ± 3 years) and all data were included in the data processing. The total scan time per volunteer was 100 minutes and the measurement was well tolerated by all volunteers.

### Data acquisition

High-resolution 2D FLASH images (field-of-view: 192x192 mm^2^, in-plane resolution: 0.6x0.6 mm^2^, slice thickness: 3 mm, 25 slices, TE/TR 9/378 ms, flip angle: 25°, acquisition time: 2:33 min) were acquired in the sagittal and transversal directions to guide voxel placement in the occipital cortex. Before the spectroscopy measurements, static magnetic B_0_ shimming was performed using the Siemens second-order shimming method.

For spatial localization, a stimulated echo acquisition mode (STEAM^25^) sequence (TM/TR 5/5000 ms) optimized for phosphorus spectroscopy in the human brain at 9.4T was used, similar as recently described^24^. Hamming-windowed sinc excitation pulses with a flip angle of about 90°, estimated from phantom B_1_ ^+^ maps, and a pulse duration of 1.5 ms were used for slice selection. The time-bandwidth product was set to 6.0 (corresponding to six zero-crossings of the amplitude modulation) resulting in an excitation bandwidth of 4.3 kHz. Thus, the chemical shift displacement error (CSDE) was 3.8% per ppm in each voxel dimension. Spoiler gradients and phase cycling scheme were optimized using DOTCOPS^24,27^. Spoiler gradients were arranged as shown in Dorst et al.^24^ with a spoiling moment of 18 ms·mT/m after the first and third slice selective pulse, and 36 ms·mT/m during the mixing time. For phase cycling, a four-step COG(0,1,2;3)^27,28^ scheme was implemented, which, together with the spoilers, removed all unwanted coherence pathways.

T_2_ relaxation time measurements were performed by acquiring a TE series with roughly exponentially spaced TEs of 6, 8, 11, 15, 20, 30, 50, 80, 150 ms. For all spectra, a voxel of 5x5x5 cm^3^ was placed in the occipital lobe. For each TE time, spectra with 120 averages were acquired with 4096 complex sampling points, an acquisition bandwidth of 10 kHz and a measurement time of 10 minutes. The first 4 averages were taken as preparation scans and omitted from the analysis. The remaining scans were assumed to be at steady-state magnetization.

### Data preprocessing

Raw data were reconstructed with an in-house written MATLAB software. The processing steps comprise averaging, singular value decomposition coil combination based on PCr^29^ (weights were calculated from filtered data and applied to non-filtered data), zero-order phase correction, and aligning PCr to 0 ppm.

The zero-order phase was calculated by maximizing the amplitude as well as the integral of the real part of PCr in the frequency domain and calculating its mean. No first-order phase correction was needed.

The full width at half maximum (FWHM) was calculated by increasing the sampling rate by a factor of 20 and finding the maximum peak height of each metabolite in a search area of 0.2 ppm around literature values for each metabolite peak frequency as well as the FWHM using an in-house written MATLAB function.

The SNR was calculated as the ratio between the metabolite peak heights and the spectral noise between +15 ppm and +30 ppm in the real part of the spectrum.

### Spectral fitting

Spectra were fitted with LCModel (version 6.3-1L)^20^ using basis sets simulated in Vespa (v1.0.0)^21^. In Vespa, a density matrix-based spectral simulation employing RF pulse waveforms in agreement with experimentally used realistic shapes and timings was performed for each metabolite and all the TEs specified. The following metabolites were simulated: phosphoethanolamine and phosphocholine (PE, PC), extracellular and intracellular free inorganic phosphate (P_*i*_^ext^, P_*i*_^int^), glycerophosphoethanolamine and glycerophosphocholine (GPE, GPC), phosphocreatine (PCr), γ- and α-adenosine triphosphate (γ-ATP, α- ATP), and nicotinamide adenine dinucleotide (NAD+, NADH). For the simulation, published chemical shifts relative to PCr at 0 ppm and homonuclear and heteronuclear J-coupling constants were used, as summarized in Table 1. γ-ATP and α-ATP moieties were simulated separately to account for possible different relaxation times of the moieties. The Vespa simulated metabolite spectra were then imported into MATLAB where the linewidth of each metabolite spectrum was adjusted to a Lorentzian linewidth similar to in vivo linewidths (see Table 2), the peaks were phase corrected, and the absolute of the peak integral was normalized. An artificial reference peak needed in the basis spectra for referencing was added at 15 ppm before the creation of LCModel basis sets.

**Table 1:** Literature phosphorus chemical shifts and scalar coupling constants used to simulate ^31^P basis spectra in Vespa.

| Metabolite | Chemical Shift [ppm] | J<sub>PP</sub>-coupling [Hz] | J<sub>PH</sub>-coupling [Hz] |
| --- | --- | --- | --- |
| PCr | 0 |  |  |
| α-ATP | -7.56<sup>30</sup> | J<sub>Pα-Pβ</sub> = 16.3<sup>30,32</sup> | J<sub>Pα-H5</sub> = 6.5<sup>1,30,33</sup><br>J<sub>Pα-H5'</sub> = 4.9<sup>1,30,33</sup><br>J<sub>Pα-H4</sub> = 1.9<sup>1,30,33</sup> |
| β-ATP | -16.18<sup>30</sup> | J<sub>Pβ-Pγ</sub> = 16.1<sup>32</sup> |  |
| γ-ATP | -2.53<sup>30</sup> |  |  |
| P<sub>i</sub><sup>int</sup> | 4.82<sup>9</sup> |  |  |
| P<sub>i</sub><sup>ext</sup> | 5.24<sup>9</sup> |  |  |
| PC | 6.23<sup>30</sup> |  | J<sub>P-H1</sub> = 6.298<sup>33</sup><br>J<sub>P-H1'</sub> = 6.249<sup>33</sup> |
| PE | 6.77<sup>30</sup> |  | J<sub>P-H1</sub> = 7.288<sup>33</sup><br>J<sub>P-H1'</sub> = 7.088<sup>33</sup> |
| GPC | 2.94<sup>30</sup> |  | J<sub>P-H3</sub> = J<sub>P-H3'</sub> = 6.03<sup>33</sup><br>J<sub>P-H7</sub> = J<sub>P-H7'</sub> = 6.03<sup>33</sup> |
| GPE | 3.49<sup>30</sup> |  | J<sub>P-H3</sub> = J<sub>P-H3'</sub> = 6.23<sup>30</sup><br>J<sub>P-H7</sub> = J<sub>P-H7'</sub> = 6.23<sup>30</sup> |
| NADH | -8.1<sup>34</sup> |  |  |
| NAD+ | -8.15<sup>34,35</sup> (ribose adenine) | J<sub>P1-P2</sub> = 20.03<sup>36</sup> | J<sub>P2-H5'</sub> = 4.8<sup>35</sup><br>J<sub>P2-H5''</sub> = 5.3<sup>35</sup><br>J<sub>P2-H4'</sub> = 2.2<sup>35</sup> |
|  | -8.45<sup>34,35</sup> (ribose nicotinamide) |  | J<sub>P1-H5</sub> = 4.3<sup>35</sup><br>J<sub>P1-H5'</sub> = 5.5<sup>35</sup><br>J<sub>P1-H4</sub> = 2.9<sup>35</sup> |

**Table 2:**
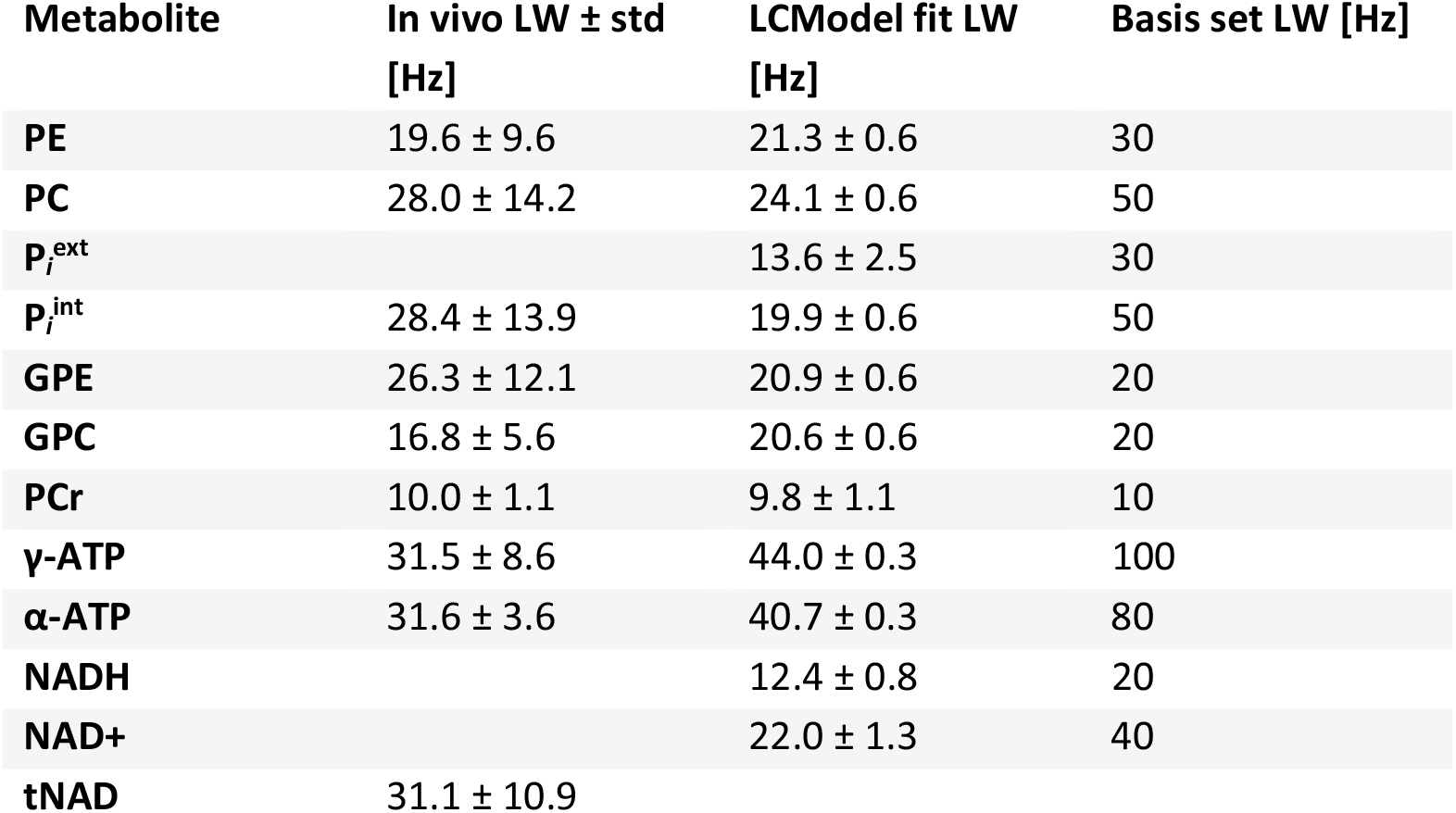
Mean spectral linewidths (LW) and their standard deviations extracted from processed raw data and LCModel fits at TE = 6 ms, as well as LWs used for the basis sets in LCModel. The simulated metabolite basis sets were line broadened to compensate large differences in T_2_* relaxation times and pH effects to best fit the spectra across the TE series.

| Metabolite | In vivo LW ± std [Hz] | LCModel fit LW [Hz] | Basis set LW [Hz] |
| --- | --- | --- | --- |
| PE | $19.6 \pm 9.6$ | $21.3 \pm 0.6$ | 30 |
| PC | $28.0 \pm 14.2$ | $24.1 \pm 0.6$ | 50 |
| $P_i^{\text{ext}}$ | | $13.6 \pm 2.5$ | 30 |
| $P_i^{\text{int}}$ | $28.4 \pm 13.9$ | $19.9 \pm 0.6$ | 50 |
| GPE | $26.3 \pm 12.1$ | $20.9 \pm 0.6$ | 20 |
| GPC | $16.8 \pm 5.6$ | $20.6 \pm 0.6$ | 20 |
| PCr | $10.0 \pm 1.1$ | $9.8 \pm 1.1$ | 10 |
| γ-ATP | $31.5 \pm 8.6$ | $44.0 \pm 0.3$ | 100 |
| α-ATP | $31.6 \pm 3.6$ | $40.7 \pm 0.3$ | 80 |
| NADH | | $12.4 \pm 0.8$ | 20 |
| NAD+ | | $22.0 \pm 1.3$ | 40 |
| tNAD | $31.1 \pm 10.9$ | | |

To perform LCModel analyses for ^31^P spectra, several adjustments are required, as described in Deelchand et al.^30^ In comparison to adjustments reported in his publication, we set the standard deviation of the first-order phases SDDEGP = 0 to not allow first-order phase correction. In addition, SDSH control parameters, which specify the standard deviation of the chemical shift, were adjusted for P_*i*_^int^, P_*i*_^ext^, γ-ATP and α-ATP to account for pH dependent frequency shifts^31^. To display correct x-axis frequencies with PCr set to 0 ppm, control parameters were adjusted to SHIFMX(2) = -4.63 and SHIFMN(2) =-4.77 defining the range about the expected value of the referencing shift^31^. LCModel analyses were then performed over the spectral range from -10 ppm to 10 ppm.

### T_2_ calculation

To calculate the apparent T_2_ relaxation times, the LCModel-fitted metabolite concentrations were fit to a single exponential two-parameter decay across the TE series according to

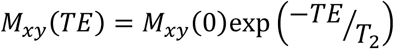

with M_xy_ being the transverse magnetization. The goodness of the fit statistics was evaluated by the coefficients of determination R^2^. T_2_ estimates with R^2^ < 0.5 were discarded from further analyses.

### pH estimation

The chemical shift difference between PCr (0 ppm) and free phosphate (*δ* in ppm) was used to calculate pH values from the modified Henderson-Hasselbalch equation^9^

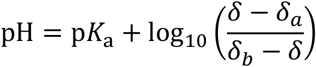

where p*K*_a_ = 6.73 is the acid dissociation constant, and *δ*_a_ = 3.275 ppm and *δ*_b_ = 5.685 ppm the chemical shifts of the protonated and deprotonated forms of free phosphate, respectively^9^. Intracellular as well as extracellular pH values were calculated from LCModel-fitted frequencies of P_*i*_^int^ and P_*i*_^ext^.

### Metabolite quantification

For metabolite quantification, peak areas S(RD, TE, TM) obtained from LCModel fits were corrected for relaxation losses according to

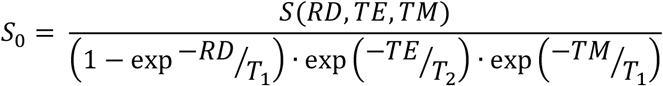

with *RD* = *TR* - *TM* - *TE*/2 ^37^. TM is the mixing time of the STEAM sequence, TE is the echo time, T_1_ are the metabolite specific longitudinal relaxation times taken from Pohmann et al.^38^, and T_2_ are the metabolite specific transversal relaxation times calculated from summed spectra in this paper (Table 3). The relaxation corrected peak areas were converted to ^31^P metabolite concentrations using an assumed γ-ATP concentration of 3 mM as an internal reference^9,39,40^.

**Table 3:**
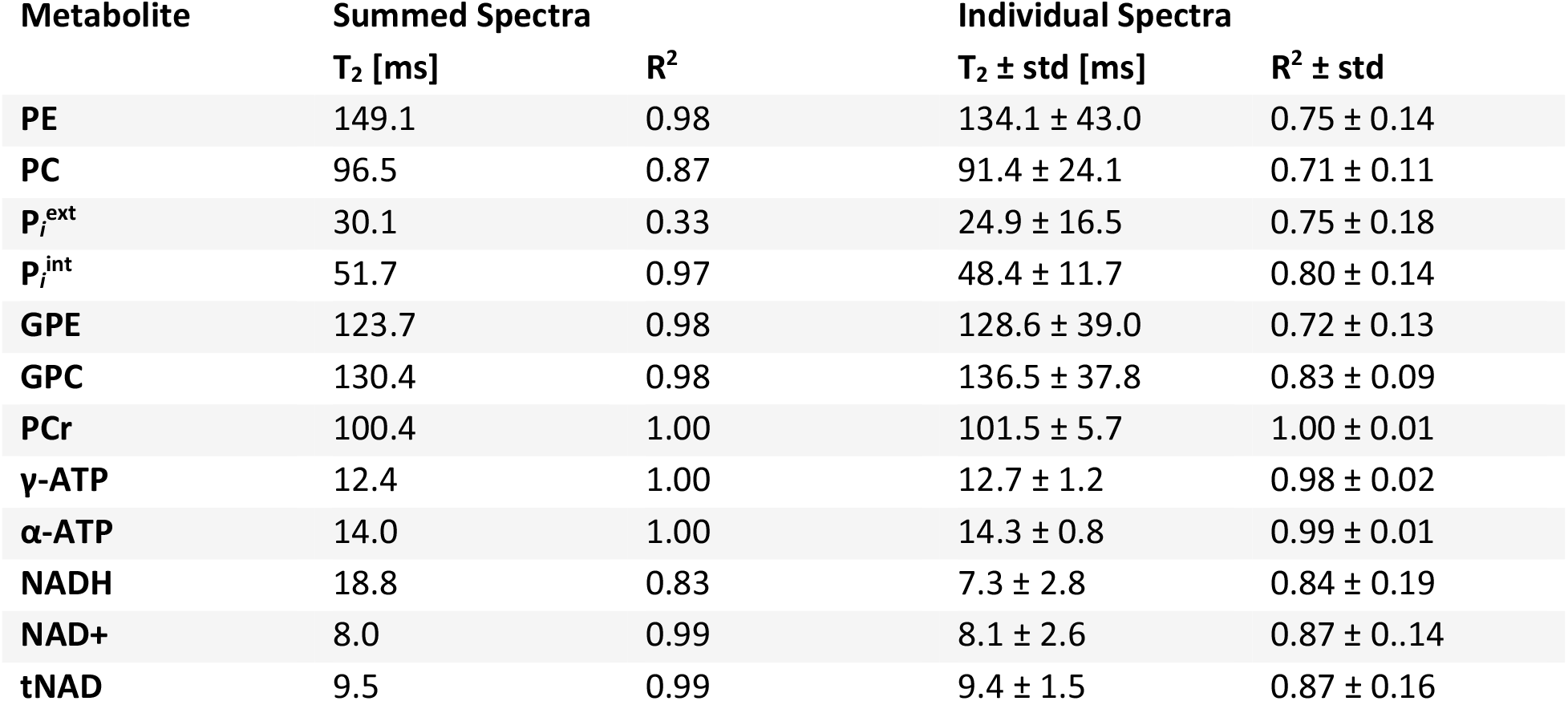
Apparent ^31^P T_2_ relaxation times in the human brain and coefficients of determination R^2^ for the across volunteers summed spectra (n = 12) as well as mean values and standard deviations derived from the individual spectra.

## Results

High-quality spectra were obtained for all volunteers and all acquired TEs with a mean PCr SNR of 36.1 ± 5.9 at TE 6 ms, and of 9.1 ± 2.7 at TE 150 ms, and similar spectral quality at TE 6 ms and TE 150 ms of 10.0 ± 1.1 Hz and 9.7 ± 1.6 Hz, respectively. Mean spectra across all volunteers and their standard deviations are shown for the echo time series in Figure 1. The shaded areas represent standard deviations across all volunteers indicating high reproducibility. Figure 2 shows the LCModel fit result for the summed spectrum at TE 6 ms from -10 ppm to 8 ppm with low fit residual. Mean spectral linewidths across all volunteers and their standard deviations are reported in Table 2 for TE 6 ms for preprocessed raw data and LCModel fits. In preprocessed raw data, linewidths of NADH and NAD+ cannot be measured separately. In addition, the linewidth P_*i*_^ext^ could not be measured reliably on a single volunteer basis. In the last column, linewidths used in LCModel basis sets are reported. The simulated metabolite basis sets were line broadened, following the observation of Deelchand et al^30^. Deelchand et al. applied line broadenings based on the measured in vivo linewidths, to compensate large differences in T_2_* relaxation times and pH effects. The Lorentzian line broadening factor in this study had to be somewhat larger than the measured linewidths to best fit the spectra across the TE series.

**Figure 1:**
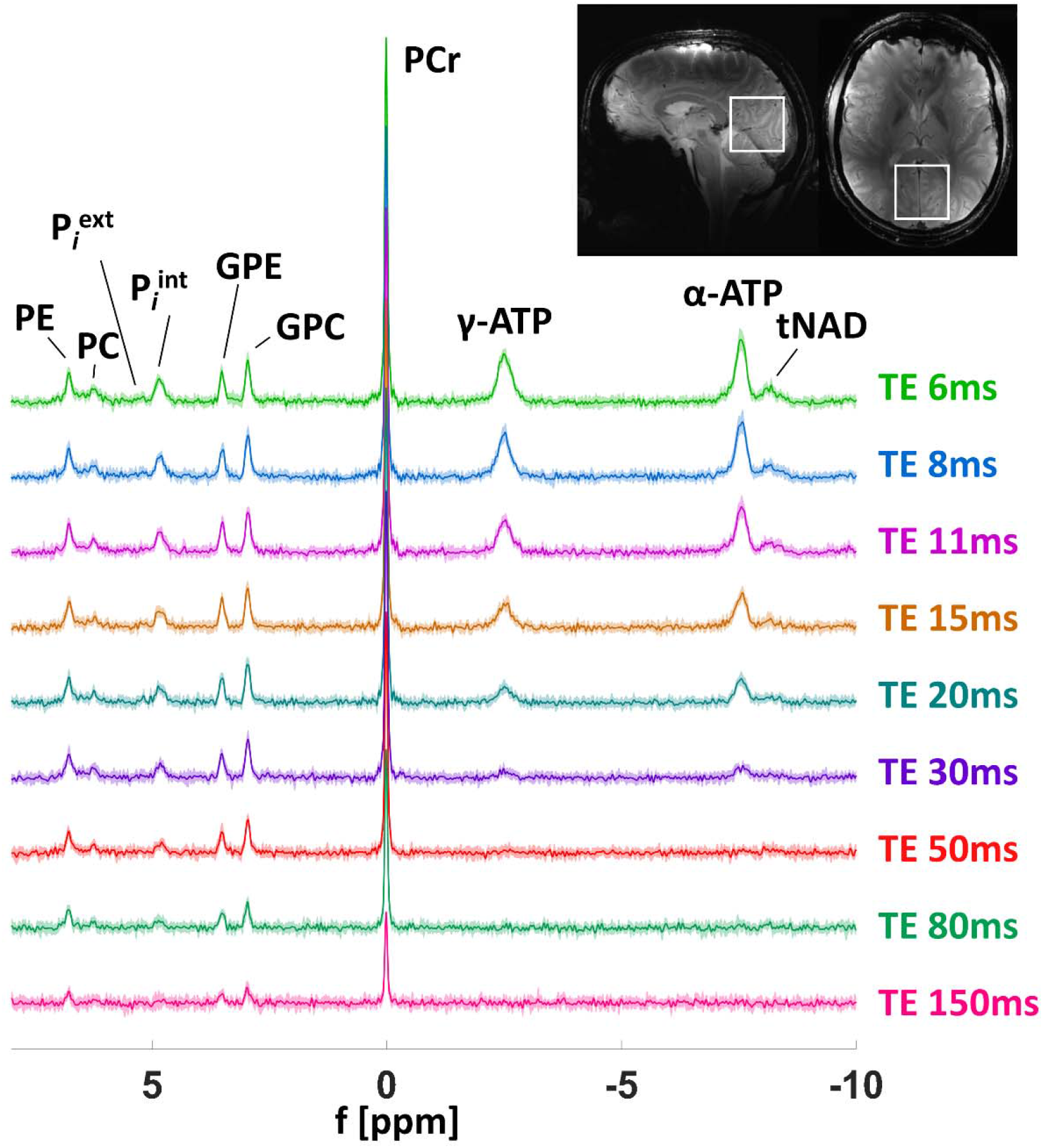
TE series (TE = 6, 8, 11, 15, 20, 30, 50, 80 and 150 ms) of the acquired ^31^P spectra. The solid line represents mean and the shaded area standard deviations across all 12 volunteers. For representation, spectra were truncated after 90 ms with subsequent zero filling back to 4096 complex sampling points. The figure insets show the voxel positioning (5x5x5 cm^3^) on acquired FLASH images in sagittal and transversal directions.

**Figure 2:**
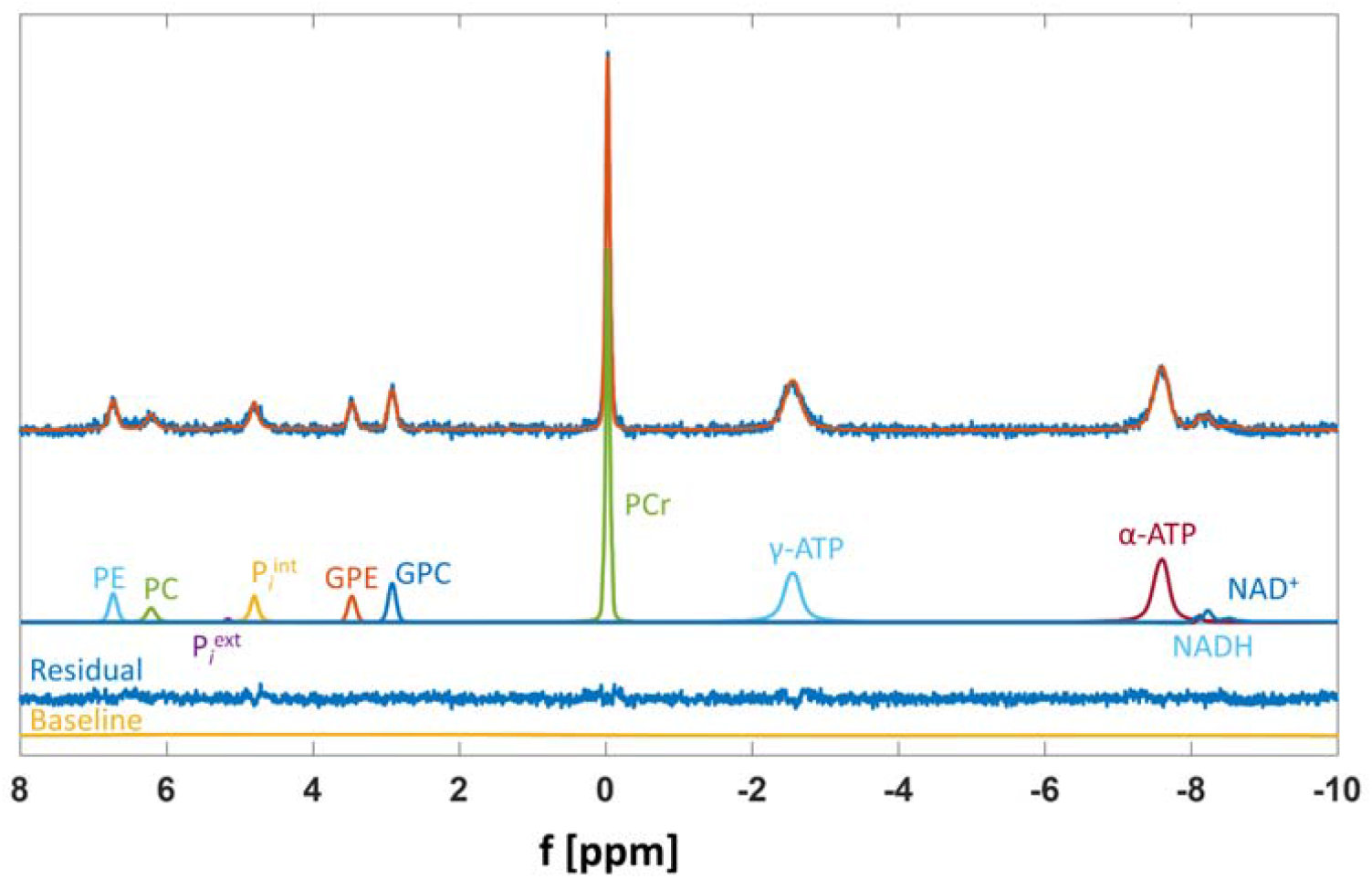
Summed spectrum averaged across all 12 volunteers with the corresponding LCModel fitted resonances at TE = 6 ms. Shown are all relevant metabolites, the corresponding fit residual and the fitted baseline. Spectra were not truncated.

Mean estimated T_2_ decay curves are presented in Figure 3 for all fitted metabolites as well as for tNAD (summed NADH and NAD+ fits). Data points and error bars represent mean fitted peak integrals over all volunteers and their standard deviations in arbitrary units. The calculated apparent T_2_ relaxation times of summed spectra as well as the mean over all volunteers (after exclusion of R^2^ < 0.5) are reported in Table 3 together with the corresponding coefficients of determination R^2^. Mean coefficients of determination of individual spectra are higher than 0.71, and of summed spectra higher than 0.83 except for P_*i*_^ext^ showing the goodness of the T_2_ fits to individual datasets. The calculated T_2_ relaxation times are visualized in boxplots in decreasing order in Figure 4 and span a wide range between ∼150 ms and ∼7 ms. A literature comparison of T_2_ relaxation times in the human brain at different field strengths is presented in Table 4.

**Table 4:** Literature T_2_ relaxation times in the human brain at various field strengths. At lower field strength, phosphomono- and diesters (PME, PDE) could not be fitted separately and are reported as one T_2_ value. * PDE was fit as a biexponential function resulting in two T_2_ values. ** Phase modulation was not taken into account

| Metabolite | Jung et al.,<br>1.5T<sup>13</sup> | Lara et al.,<br>2T<sup>14</sup> | Merboldt et<br>al., 2T<sup>15</sup> | Lei et al.,<br>7T<sup>11</sup> | Van der Kemp<br>et al., 7T<sup>16</sup> |
| --- | --- | --- | --- | --- | --- |
| PME | 196 | 33 | 70 |  |  |
| PE |  |  |  |  | 202 ± 6 |
| PC |  |  |  |  | 129 ± 6 |
| Pi | 129 | 81 | 80 |  | 86 ± 2 |
| PDE | 3 / 253 * | 11 | 20 |  |  |
| GPE |  |  |  |  | 214 ± 10 |
| GPC |  |  |  |  | 213 ± 11 |
| PCr | 241 | 390 | 150 | 132.0 ± 12.8 |  |
| γ-ATP | 89 ± 9 | 17** | 30** | 26.1 ± 9.6 |  |
| α-ATP | 84 ± 6 | 28** | 30** | 25.8 ± 6.6 |  |
| β-ATP | 62 ± 3 | 15** | 20** |  |  |

**Figure 3:**
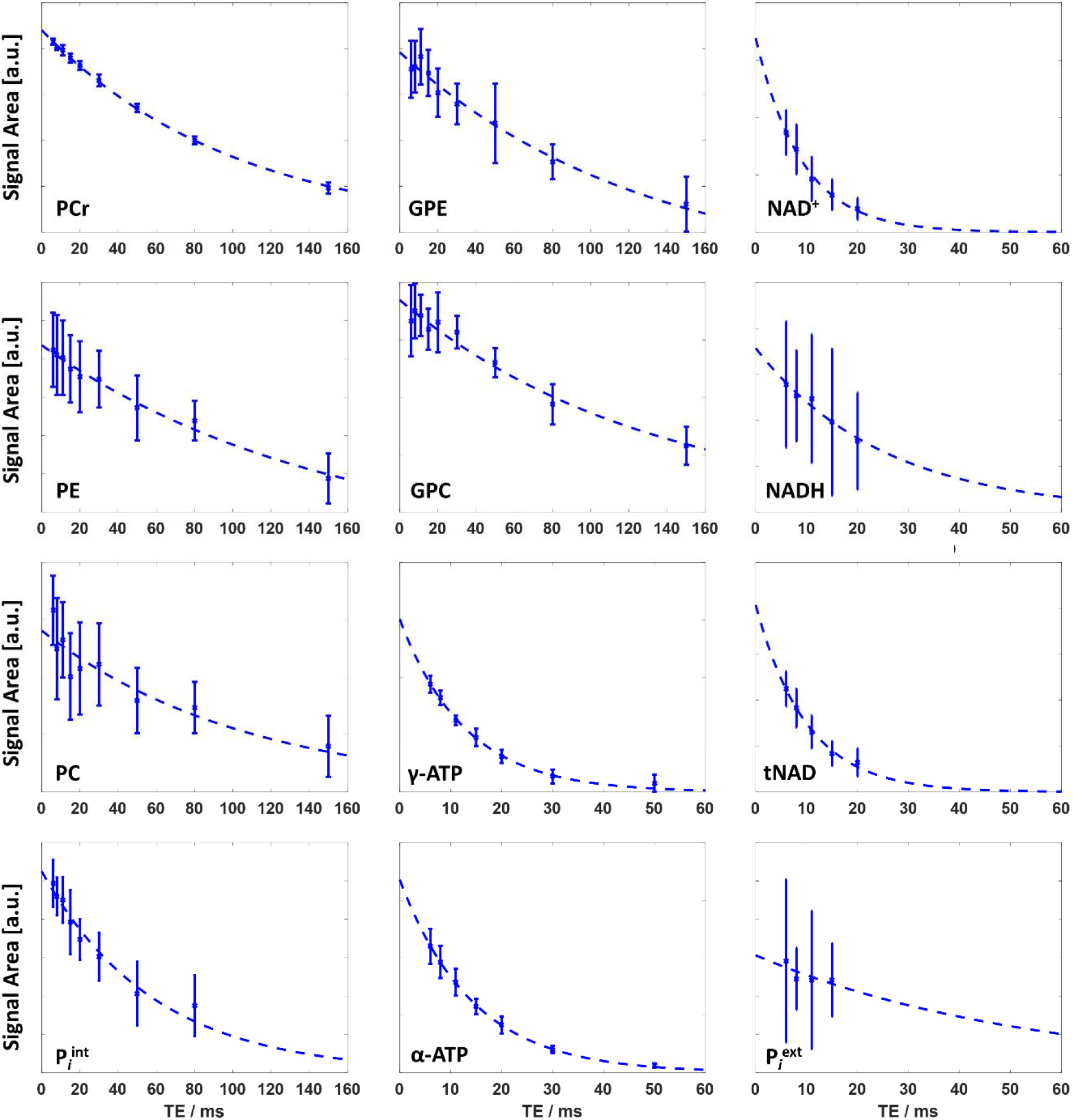
Mean estimated T_2_ decay curves of all 11 fitted metabolite peaks and summed tNAD (dashed lines). The error bars show the standard deviations of the fitted metabolite peak integrals across all 12 volunteers in arbitrary units. Apparent T_2_ times and coefficients of determination R^2^ are listed in Table 3.

**Figure 4:**
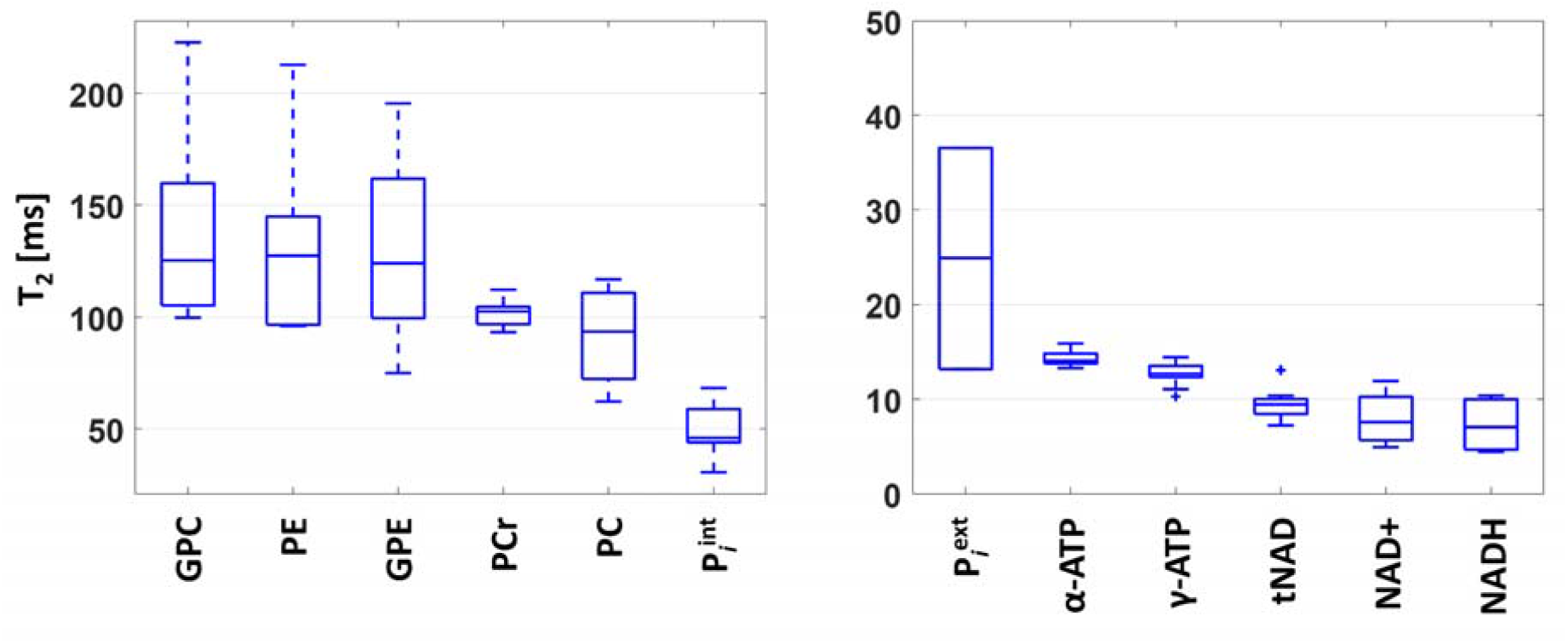
Estimated apparent T_2_ relaxation times in ms calculated from nine TE times across 12 volunteers. The central mark in each box indicates the median; the bottom and top edges denote the 25^th^ and 75^th^ percentiles, respectively. The error bars correspond to the standard deviation over all measured volunteers. Apparent T_2_ times and coefficients of determination R^2^ are listed in Table 3.

Relaxation-corrected LCModel quantification results for eleven ^31^P resonances and tNAD acquired from spectra of 12 healthy volunteers with TE 6 ms are presented in Figure 5. Concentration values are given in mM with reference to γ-ATP as an internal reference. For P_*i*_^ext^, no T_1_ correction could be applied; for NADH, NAD+ and tNAD, the T_1_ relaxation time reported for tNAD was used^38^. Relaxation-corrected, as well as non-corrected concentrations, are also reported in Table 5 calculated from summed as well as individual spectra.

**Table 5:** Fit concentrations in arbitrary units and relaxation-corrected quantification results in mM for eleven ^31^P resonances and tNAD for summed spectra as well as individually fit spectra at TE 6 ms. Additionally, pH values are reported. Errors of the concentrations and pH values calculated from individual volunteers are their corresponding standard deviations. *: for extracellular P_i_, no T_1_ correction could be applied. **: for NADH, NAD+ and tNAD the same T_1_ relaxation time was assumed.

| Metabolite | LCModel concentrations [a.u.] | Summed Spectra, $T_1$&$T_2$ corr [mM] | Individual Spectra, $T_1$&$T_2$ corr [mM] |
| --- | --- | --- | --- |
| PE | $0.86 \pm 0.26$ | 0.85 | $0.85 \pm 0.21$ |
| PC | $0.51 \pm 0.13$ | 0.39 | $0.40 \pm 0.09$ |
| $P_i^{\text{ext}}$ * | $0.13 \pm 0.13$ | 0.06 | $0.20 \pm 0.14$ |
| $P_i^{\text{int}}$ | $0.80 \pm 0.19$ | 0.70 | $0.68 \pm 0.10$ |
| GPE | $0.74 \pm 0.18$ | 0.66 | $0.66 \pm 0.12$ |
| GPC | $1.09 \pm 0.28$ | 0.99 | $0.97 \pm 0.21$ |
| PCr | $6.69 \pm 1.15$ | 5.07 | $4.88 \pm 0.34$ |
| γ-ATP | $3.00 \pm 0.41$ | 3.00 | $3.00 \pm 0.00$ |
| α-ATP | $3.48 \pm 0.37$ | 3.19 | $3.23 \pm 0.24$ |
| NADH ** | $0.16 \pm 0.08$ | 0.12 | $0.21 \pm 0.03$ |
| NAD$^{+}$ ** | $0.56 \pm 0.14$ | 0.76 | $0.73 \pm 0.15$ |
| tNAD ** | $0.71 \pm 0.13$ | 0.84 | $0.84 \pm 0.13$ |
| pH |  |  |  |
| pH$_{\text{int}}$ | | 6.99 | $6.98 \pm 0.02$ |
| pH$_{\text{ext}}$ | | 7.32 | $7.30 \pm 0.18$ |

**Figure 5:**
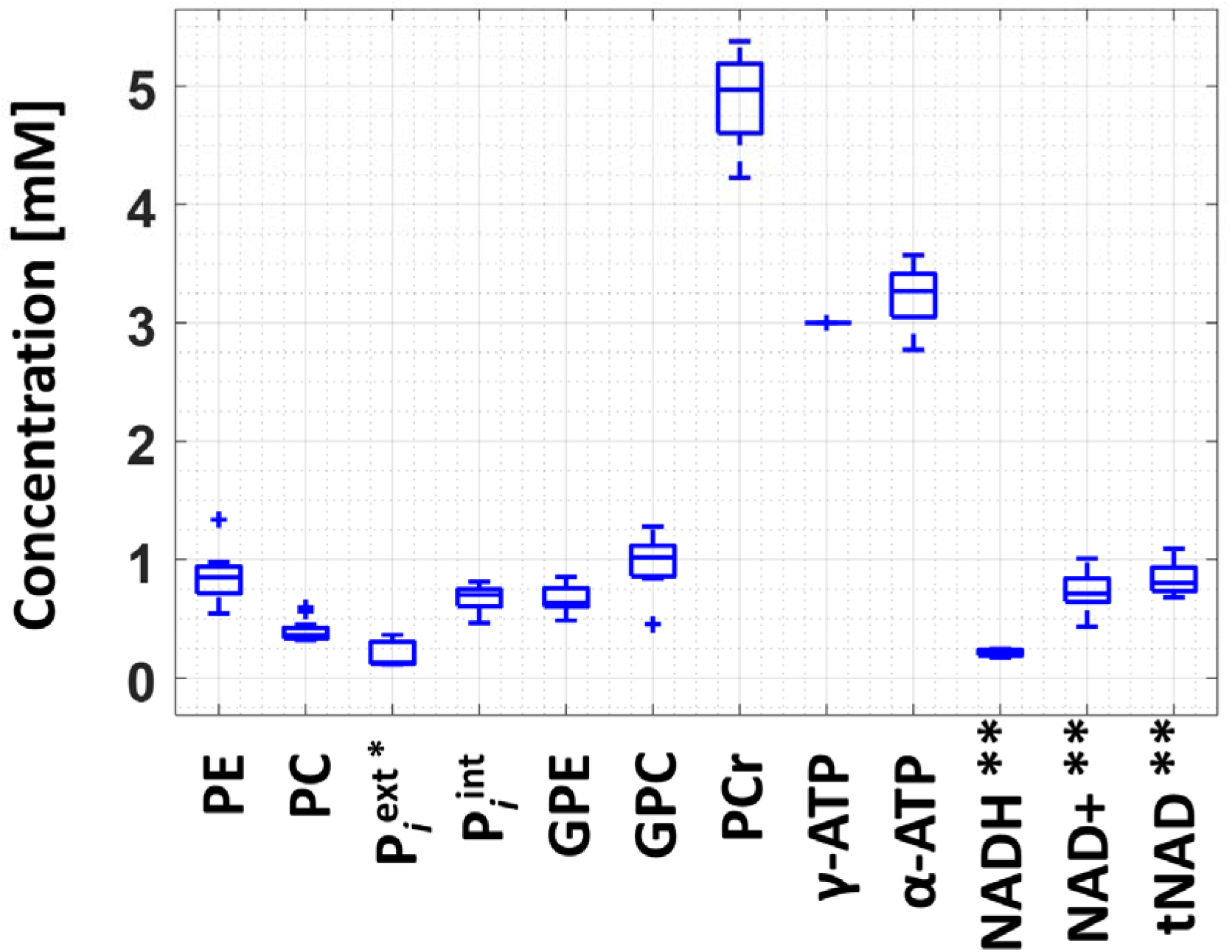
Relaxation-corrected LCModel quantification results from spectra acquired from 12 healthy volunteers. Concentration values are given in mM with reference to γ-ATP as an internal reference. The central mark in each box indicates the median; the bottom and top edges denote the 25th and 75th percentiles, respectively. The error bars correspond to the standard deviation over all measured volunteers. * For P_i_^ext^, no T_1_ correction could be applied. ** for NADH, NAD+ and tNAD, the same T_1_ relaxation time was assumed.

## Discussion

This study presents in vivo localized T_2_ relaxation times of singlets as well as J-coupled spin resonances of human cerebral metabolites detectable with ^31^P MRS at 9.4T. The high spectral resolution in combination with accurately simulated basis sets allowed the estimation of 10 metabolites T_2_s acquired in one TE spectra series, including for the first time T_2_s of NAD+ and NADH measured with ^31^P MRS and P_*i*_^ext^. Also, metabolite concentrations, as well as intracellular and extracellular pH, were calculated.

### Spectral quality

Even though the localization efficiency of STEAM is lower than for adiabatic sequences, the single-shot STEAM sequence offers higher SNR per unit time and allows for very short TEs, which are both critical factors for the acquisition of metabolite signals with short relaxation times^24^. The spectra obtained in this study with a STEAM sequence allowing a minimum TE of 6 ms show good SNR for a measurement time of 10 minutes. The transmit frequency was set to PCr to cover most ^31^P metabolites. With this setup, β-ATP at -16.26 ppm could not be measured due to the limited bandwidth of the excitation pulses and the high chemical shift displacement error. SNR of the visible metabolite peaks decays exponentially with increasing TE with some peaks still visible at TE 150 ms.

### T_2_ estimation methods

The high spectral quality allowed spectral fitting on a single volunteer basis. Based on the quality of the LCModel fit and residual, as shown in Figure 2, the basis sets generated in Vespa using chemical shifts and J-coupling constants given in Table 1 appear suitable for analyzing ^31^P MRS data. Since J-evolution was considered in the basis sets, measurement of spectra at any echo time desired is possible.

With this approach, neither frequency-selective refocusing or homonuclear decoupling^11,13,16,18,19,23^ nor assumptions of signal loss at specific TE of phase-modulated metabolites^41^ nor measurement at specific TEs to exactly match multiples of 1/J to completely refocus the metabolite of interest^17,18^ are necessary. In a frequency-selective spin-echo method, the frequency-selective pulse only affects one of the coupling nuclei (γ- and α-ATP or β-ATP), while the other peaks are presented without distortions^18^. Therefore, no assumptions about J-coupling constants are necessary and spectra at any TE can be acquired for T_2_ measurements. However, when using frequency selection, two series of acquisitions are necessary to obtain spectra of all three ATP nuclei resulting in double the measurement time. When using selective spin decoupling, the effects of phase modulation are suppressed by selectively irradiating one of the coupling nuclei during the period of signal acquisition. The difficulty is to selectively irradiate a single resonance to avoid saturation spillover onto neighboring resonances. To ensure complete decoupling, the decoupler power needs to be sufficiently high. This results in higher frequency spread and might disturb resonances close in frequency to the target^19^. Another option to measure T_2_ without sequence modifications is to measure undistorted signals by choosing TE a multiple of 1/J. However, for the three different ATP moieties, this is fulfilled at different echo times since their J-coupling constants are slightly different^17,18^.

### T_2_ relaxation

The method chosen in this study to acquire and fit data with accurately simulated spectral basis sets comprising J-evolution resulted in reliable estimation of T_2_ relaxation times on a single volunteer basis for 10 ^31^P metabolites. For P_*i*_^int^, PCr, γ- and α-ATP, which are involved in chemical exchange and/or cross-relaxation, the measured dephasing rates reflect the apparent T_2_ relaxation times. The T_2_ relaxation times of all measured metabolites could be reliably estimated in individual volunteers as well as from spectra summed across all volunteers with coefficients of determination higher than 0.7 except for P_*i*_^ext^.

Leaving the two studies at 2T, where the influence of the homonuclear J-coupling of ATP on the behavior of spin echoes was not completely accounted for, aside^14,15^, our results fit into a decreasing trend of T_2_ relaxation times with increasing field strength B_0_ ^11,13,16^. This is in line with the theory of the two competing mechanisms of dipolar proton-phosphorus interactions and chemical shift anisotropy (CSA) considered to determine ^31^P relaxation times^1,23^. While the influence of dipolar relaxation decreases with increasing main magnetic field strength, CSA relaxation increases with the square of the main magnetic field strength^1,23,42^. It is therefore expected that CSA becomes increasingly important for the relaxation mechanisms in ^31^P metabolites with increasing B_0_ ^43^. The relative contribution of each mechanism to the relaxation times also depends on the phosphate group present in different metabolites^43^. Since relaxation as a result of CSA correlates with the symmetry of the magnetic shielding, a stronger decrease in T_2_ with increasing B_0_ is expected for less symmetric molecules^1,23^.

The much shorter T_2_ relaxation times of ATP in comparison to the other ^31^P MRS detectable metabolites are in accordance with literature values where J-evolution was taken into account^11,13,16^. This has been suggested to possibly originate partly from the exchange between bound and free states of ATP, its interaction with creatine kinase and ATPase enzymes, and its strong interaction with the unpaired electron of complexed paramagnetic ions besides CSA^1^. A further possible explanation for the short relaxation times is that there might be a strong ^31^P dipole-dipole interaction through the bond –P-O-P-, which is also true for NAD+ and NADH^9^. Measured T_2_ relaxation times of NAD+ and NADH are also very short below 20 ms and in a similar range as estimated ATP T_2_ relaxation times. Estimated NAD+ and NADH T_2_ relaxation times are similar, which is in accordance with assumptions made in ^1^H measurements^35^.

The estimated T_2_ relaxation times of PE, GPE and GPC are the longest and similar at approximately 130 ms. The estimated T_2_ of PC is lower at approximately 95 ms. This was also seen in a 7T study^16^ and attributed to a contribution of 2,3-diphosphoglycerate (2,3-DPG) from blood to the signal of PC, but to a lesser extend to PE. It was shown that when accounting for 2,3-DPG, which has a shorter apparent T_2_ than PC, the estimated T_2_ of PC becomes similar to T_2_s of PE, GPE and GPC.

Estimated apparent T_2_s of inorganic phosphate are rather short in comparison to T_2_s of phosphomono- and –diesters. The short apparent P_*i*_^int^ could be explained by the exchange of P_*i*_^int^ with γ-ATP via glycolysis which has short relaxation times^9,16,39^. However, the T_2_ of P_*i*_^ext^ measured in our study is lower than of P_*i*_^int^, although the extracellular P_i_ pool was shown to be metabolically inactive^9,16,44^. Yet, it has to be mentioned that P_*i*_^ext^ could not be fitted in every volunteer reliably and its SNR is low which results in an uncertainty of the estimated T_2_ relaxation time, reflected in the low coefficient of determination R^2^ of P_*i*_^ext^ in the summed spectra analysis.

The shorter apparent T_2_ of PCr in comparison with phosphomonoesters and –diesters can also be explained by the chemical exchange of PCr with γ-ATP^9,39^. Additionally, the chemical surrounding of the phosphorus atom (bond to three oxygen and one nitrogen atom) might attribute to a relatively large contribution of CSA to relaxation in comparison to the phosphomonoesters and –diesters (P atoms bond to four oxygen atoms) resulting in faster relaxation of PCr^9^.

While the apparent T_2_ relaxation times of metabolites with high SNR could be estimated reliably, the signal intensity tends to be overestimated with increasing TE and, therefore, decreasing SNR^2^. This might induce a systematic error in T_2_ estimation, which could be compensated by acquiring more averages for longer TEs. However, acquiring more averages results in longer scan durations not feasible in this study. Also, the assumption about J-modulation used in this study to model basis sets could influence the reliability of estimated T_2_ in coupled spin systems. Even though our results fit in a decreasing trend of T_2_ with increasing B_0_, a more detailed comparison to published data at lower field strengths is difficult due to the application of different measurement methods and hardware setups, different data processing and fitting, and different volunteer populations measured.

### pH estimation

The estimated intracellular and extracellular pH values were observed to be consistent among the two different fitting types. Compared to earlier studies, our estimated values of intracellular and extracellular pH agree well with pH_int_ 6.98 and pH_ext_ 7.32 measured in the human brain at 7T with a pulse-acquire sequence in combination with 4 outer volume suppression bands^45^. Our pH_int_ and pH_ext_ values also confirm those measured with 3D ^31^P MRSI at 7T^4^ and 9.4T^46^ in mixed grey and white matter tissue.

### Metabolite concentrations

The majority of metabolite concentrations obtained in this study are comparable to values reported in Ren et al.^9^ measured at 7T using a pulse-acquire sequence and a long TR of 25 s to guarantee full T_1_ recovery. The tNAD concentration measured in this study is higher than the one measured in Ren et al.^9^, whereas the PE and P_*i*_^ext^ concentrations from this study are much lower. Since uridine diphosphoglucose (UDPG), which has overlapping resonance frequencies with NAD+ and NADH, was not fitted in this study, the obtained tNAD concentration might be influenced by UDPG^34,47^. Furthermore, the low abundance of NADH and NAD+ and the resulting low SNR might have induced fit errors and errors in their T_2_ estimation. The much lower P_*i*_^ext^ concentration might result from different measurement techniques. In comparison to the 7T study, localized spectra were acquired in this study. As was shown in a previous study, the extracellular P_i_ peak might originate from blood and CSF in the peripheral region of the brain^45^. This explanation is supported by a previously published 3D MRSI study reporting similar P_*i*_^ext^ concentrations to this study^46^. However, it is also possible that the P_*i*_^ext^ concentration reported in this study is slightly too low due to too low estimated T_2_ relaxation times as a result of possible muscle tissue influence on our data. Although the PCr concentration calculated in this study fits to Ren et al.^9^, it is slightly higher than concentrations calculated in Ruhm at al.^46^, employing 3D MRSI at 9.4T. This also might hint at small contamination of brain spectra by muscle tissue, since the PCr concentration in muscle tissue is much higher than in the brain^40,48^. The metabolite concentrations of all metabolites obtained in this study are about 40% lower than in a recently published ISIS localized study at 9.4T, except for ATP, tNAD and PCr ^24^. Although ISIS localization implies a short TE, fitting simple peak integrals to analyze multiplets that undergo J-evolution is not sufficient to determine metabolite concentrations^49^. Therefore, a possibly too low γ-ATP concentration was determined in the ISIS study, which results in too high concentrations of the other metabolites when normalized to γ-ATP.

## Conclusions

^31^P transversal relaxation times of human brain metabolites at 9.4T are reported, including values for P_*i*_^ext^, NAD+ and NADH. To the best of our knowledge, T_2_s of ^31^P metabolites that undergo J-evolution were estimated for the first time by considering J-evolution in an accurately modeled basis set for each TE used in the fitting routine. A decreasing trend of T_2_ with increasing field strength adding to previous literature confirms the limited usability of echo-based acquisition methods for ^31^P MRS at UHFs. The estimated relaxation times were used for absolute quantification resulting in metabolite concentrations comparable to literature values measured from FIDs.

## Abbreviations used

δ: chemical shift
2,3-DPG: 2,3-diphosphoglycerate
ATP: adenosine triphosphate
COG: cogwheel phase cycling
CSA: chemical shift anisotropy
CSDE: chemical shift displacement error
FID: free induction decay
FLASH: fast low angle shot
FWHM: full-width half maximum
GPC: glycerophosphocholine
GPE: glycerophosphoethanolamine
MRS: magnetic resonance spectroscopy
NADH: nicotinamide adenine dinucleotide
NAD+: NAD oxidized
PC: phosphocholine
PCr: phosphocreatine
PE: phosphoethanolamine
pHext: extracellular pH
pHint: intracellular pH
Piext: extracellular inorganic phosphate
Piint: intracellular inorganic phosphate
R2: coefficient of determination
SAR: specific absorption rate
SNR: signal to noise ratio
STEAM: stimulated echo acquisition mode
T1: longitudinal relaxation time
T2: transversal relaxation time
TE: echo time
TM: mixing time
tNAD: total NAD
TR: repetition time
UDPG: uridine diphosphoglucose
UHF: ultrahigh-field

